# Genomic region associated with pod color variation in pea (*Pisum sativum*)

**DOI:** 10.1101/2020.09.25.313072

**Authors:** Kenta Shirasawa, Kazuhiro Sasaki, Hideki Hirakawa, Sachiko Isobe

**Author notes:** Corresponding author: Kenta Shirasawa. Japan International Research Center for Agricultural Sciences, Tsukuba, Ibaraki 305-8686, Japan (Kazuhiro Sasaki).

## Abstract

Pea (*Pisum sativum*) was chosen as the research material by Gregor Mendel to discover the laws of inheritance. Out of seven traits studied by Mendel, genes controlling three traits including pod shape, pod color, and flower position have not been identified to date. With the aim to identify the genomic region controlling pod color, we determined the genome sequence of a pea line with yellow pods. Genome sequence reads obtained using a nanopore sequencing technology were assembled into 117,981 contigs that spanned 3.3 Gb in length and showed an N50 value of 51.2 kb. Using single nucleotide polymorphisms (SNPs) detected in a pea mapping population, these contigs were genetically anchored to the publicly available pseudomolecule sequences of the pea genome. Subsequent genetic and association analyses identified a genomic region responsible for pea pod color. At this genomic location, genes encoding 3’ exoribonucleases were selected as potential candidates controlling pod color, based on DNA sequencing and transcriptome analysis of green and yellow pod lines. The results presented in this study are expected to accelerate pan-genome studies in pea and facilitate the identification of the gene controlling one of the traits studied by Mendel.

## INTRODUCTION

Advances in the field of genomics, owing to the development of next-generation sequencing (NGS) technologies, have enabled us to identify genes as DNA sequences in genomes (Nguyen et al., 2019). The field of classical genetics was initiated by Gregor Mendel through the discovery of the laws of inheritance in pea (*Pisum sativum*). Mendel studies seven traits of pea (Ellis et al., 2011; Reid and Ross, 2011; Von Mendel, 1866), including seed shape, seed color, flower color, pod shape, pod color, flower position, and stem height. However, so far, genes controlling only four of these traits, including seed shape (Bhattacharyya et al., 1990), seed color (Armstead et al., 2007; Sato et al., 2007), flower color (Hellens et al., 2010), and stem height (Lester et al., 1997), have been identified based on sequence variation in the pea genome. Given its diploid and autogamous nature, pea shows simple inheritance patterns, which is believed to have helped in the discovery of the laws of inheritance. However, the pea genome is large (∼4.5 Gb) (Arumuganathan and Earle, 1991; Greilhuber and Ebert, 1994) and complex because of highly repetitive sequences (Macas et al., 2007; Murray et al., 1981). These characteristics of the pea genome interfere with its genomic analysis.

To the best of our knowledge, two genome assemblies of pea are publicly available, one of which is a chromosome-level reference genome assembly, ‘Pisum_sativum_v1a’, of a French pea cultivar ‘Caméor’ (Kreplak et al., 2019), while the other is a draft genome assembly of a pea cultivar ‘Gradus No 2’ (ASM301357v1) released by the Earlham Institute, UK, under the GenBank assembly accession number GCA_003013575.1. The Pisum_sativum_v1a assembly consists of seven pseudomolecules (3.3 Gb) anchored to pea chromosomes, and 14,267 scaffold sequences (685.4 Mb) unassigned to any chromosome but potentially harboring 44,756 genes. Additionally, repetitive sequences account for ca. 83% of this genome assembly. On the other hand, the ASM301357v1 assembly (4.3 Gb) consists of as many as 5.4 million scaffold sequences; however, no annotation information is available. These two reference genome assemblies could help identify all genes and genetic variations present within *P. sativum* (Gan et al., 2011).

Pan-genomes represent core genes found in all individuals within a species, in addition to dispensable genes present in only some accessions (Bayer et al., 2020). However, repetitive sequences present in the genome might inhibit the alignment of sequence reads to the reference genome assembly, decreasing the accuracy of pan-genome studies. Instead of sequence similarity searches, genetic anchoring of multiple reference sequences with DNA markers would provide more accurate pan-genome results. NGS-based genotyping techniques (Davey et al., 2011), such as double digest restriction-site associated DNA sequencing (ddRAD-Seq; (Peterson et al., 2012), would be effective for the comparative analysis of genome sequences and structures. For instance, a set of ddRAD-Seq reads obtained from a mapping population could be aligned to each of the multiple reference sequences for detecting single nucleotide polymorphisms (SNPs), and subsequent linkage analysis of these SNPs could cluster identical SNP loci or genomic positions within a single map position (Shirasawa and Kitashiba, 2017). This approach would be useful for assigning a sequence from one line to the counterparts.

In this study, we aimed to identify the genomic region that determines the pod color in pea. The genome of a pea line with yellow pods was sequenced, and the genome assembly was anchored to the publicly available reference sequences using SNPs obtained by ddRAD-Seq as anchors. Since the ddRAD-Seq data were derived from an F2 mapping population with segregating pod color and flower color, the color loci were genetically and physically identified by genetic mapping and genome-wide association study (GWAS) approaches. Subsequent whole genome resequencing and transcriptome analyses identified candidate genes controlling pod color and flower color in pea. Thus, the results of this study not only facilitate pan-genomic studies in pea but also enable the discovery of genes controlling traits studied by Mendel.

## EXPERIMENTAL PROCEDURES

### Plant material

Two pea lines, JI4 and JI128, possessing green and yellow pods, respectively, provided by John Innes Centre (Norwich, UK) were used in this study. The two pea lines were crossed to obtain F1 seeds. The F2 plants (n = 167) were grown in an experimental field in the Institute for Sustainable Agro□ecosystem Services, Graduate School of Agricultural and Life Sciences, The University of Tokyo (35°74′N, 139°54′E)., and the color of pods was visually evaluated during pod development.

### De novo genome assembly

DNA was extracted from the leaves of JI128 plants using the DNeasy Plant Mini Kit (Qiagen, Hilden, Germany). Short-read DNA sequencing libraries were prepared according to the manufacturer’s instructions, and sequenced on DNA sequencers, NextSeq 500 (Illumina) and DNBSEQ-G400 (MGI Tech, Shenzhen, China). The genome size of the JI128 line was estimated by k-mer distribution analysis using the Jellyfish software (Marcais and Kingsford, 2011).

High-molecular weight DNA was extracted from JI128 DNA using the Genomic-tip kit (Qiagen), and a long-read sequence library was constructed using the Rapid Sequencing Kit (version SQK-RAD004) (Oxford Nanopore Technologies, Oxford, UK). The library was sequenced with the MinION using flow cell version FLO-MIN107 R9 (Oxford Nanopore Technologies). Base calling from the FAST5 files were performed using Guppy v2.3.5 (Oxford Nanopore Technologies). The long reads were assembled with wtdbg2 v2.2 (Ruan and Li, 2020), and potential sequencing errors in the contigs were corrected once using Pilon (Walker et al., 2014). The resulting genome assembly was designated as PSA_r1.0. Assembly completeness was evaluated with the embryophyta_odb10 data using Benchmarking Universal Single-Copy Orthologs (BUSCO) v3.0.2 (Simao et al., 2015).

### ddRAD-Seq analysis

A ddRAD-Seq library for the 168 F2 lines, an F1 hybrid, and both parental lines (JI4 and JI128) were prepared using two restriction enzymes, *Pst*I and *Msp*I, as described previously (Shirasawa et al., 2016). The ddRAD-Seq libraries were sequenced on the HiSeq4000 platform (Illumina). Low-quality bases were removed from the reads using PRINSEQ v0.20.4 (Schmieder and Edwards, 2011), and adapter sequences were trimmed using fastx_clipper in the FASTX-Toolkit v0.0.13 (http://hannonlab.cshl.edu/fastx_toolkit). Bowtie2 v2.2.3 (Langmead and Salzberg, 2012) was used to map the filtered reads onto three reference genome assemblies: Pisum_sativum_v1a (Kreplak et al., 2019), ASM301357v1 (GenBank accession number PUCA000000000), and PSA_r1.0 (this study). The resultant Sequence Alignment/Map (SAM) files were converted to Binary Sequence Alignment/Map (BAM) files and subjected to SNP calling using the mpileup option of SAMtools v0.1.19 (Li et al., 2009) and the view option of BCFtools. High-confidence SNPs were selected using VCFtools v0.1.12b (Danecek et al., 2011), according to the following criteria: (1) depth of coverage ≥ 5 (for each line); (2) SNP quality scores = 999 (for each locus); and (3) proportion of missing data < 0.5 (for each locus).

### Genetic map construction

Linkage analysis of high-confidence SNPs was performed with LepMap3 v0.2 (Rastas, 2017). Marker loci were classified roughly into seven linkage groups using the SeparateChromosomes2 module (logarithm of the odds [LOD] score = 19), while considering segregation distortion. Marker order and map distances were calculated using the OrderMarkers2 module. The pod color trait was mapped on the resultant genetic map with MAPMAEKR v3.0b (Lander et al., 1987).

Association mapping was performed using the general linear model implemented in TASSEL version 5.0 (Bradbury et al., 2007). Thresholds for the association were set at a significance level of 1%, after Bonferroni multiple test correction.

### Whole genome resequencing analysis

Short-read sequence libraries for JI4 and JI128 were prepared using the TruSeq DNA PCR-Free Library Kit (Illumina), according to the manufacturer’s instructions, and sequenced on the NextSeq500 system (Illumina). Low-quality bases and adapter sequences were trimmed and mapped onto the three reference genome assemblies, as described above. High-confidence SNPs were selected using VCFtools v0.1.12b (Danecek et al., 2011), according to the following criteria: (1) depth of coverage ≥ 5 (for each line); (2) SNP quality score = 999 (for each locus); and (3) proportion of missing data < 0.5 (for each locus). Effects of SNPs on gene function were predicted using SnpEff v4.3t (Cingolani et al., 2012).

### RNA-Seq analysis

Total RNA was extracted from immature pods of JI4 and JI128 using the RNeasy Mini Kit (Qiagen). The isolated total RNA was treated with RQ1 RNase-Free DNase (Promega, Madison, WI, USA) to remove contaminating genomic DNA, and RNA libraries were constructed using the TruSeq Stranded mRNA Library Kit (Illumina). The RNA libraries were sequenced on the NextSeq 500 system (Illumina) to generate 151 bp paired-end reads. Low-quality bases were removed with PRINSEQ v0.20.4 (Schmieder and Edwards, 2011), and adapter sequences were trimmed with fastx_clipper (parameter: □a AGATCGGAAGAGC) in the FASTX□Toolkit v0.0.13 (http://hannonlab.cshl.edu/fastx_toolkit). Gene expression was quantified by mapping the RNA-Seq reads onto the PSA_r1.0 assembly using HISAT2 v2.1.0 (Kim et al., 2015), followed by sequencing depth normalization to determine the number of fragments per kilobase of exon model per million mapped reads (FPKM) using StringTie v1.3.5 (Pertea et al., 2015) and Ballgown v2.14.1 (Frazee et al., 2015), as described previously (Pertea et al., 2016).

## RESULTS

### De novo genome assembly of JI128 possessing yellow pod color

The genome sequence of JI128 was determined using the Rapid Sequencing Kit (version SQK-RAD004), a long-read sequencing technology of Oxford Nanopore Technologies (Oxford, UK). First, the size of the JI128 genome was estimated as 4.5 Gb via k-mer distribution analysis of the short-read data (147 Gb) obtained from DNBSEQ-G400 (Figure 1, Table S1). The library was sequenced on 35 flow cells with the MinION, and a total of 262.1 Gb data were obtained, consisting of 33.2 million (M) reads with an N50 read length of 15.5 kb (Table S1). Potential sequencing errors were corrected, and the cleaned reads (total 12.3 M reads; 196.7 Gb; 44× genome coverage) were assembled into 117,981 contigs. BUSCO analysis indicated only 51.0% of the complete BUSCOs in the assembly. Then, sequencing errors in the assembly were corrected using Pilon, for which Illumina and MGI short reads were employed. The final assembly was 3.3 Gb in length, with an N50 value of 51.2 kb; this assembly was designated as PSA_r1.0 (Table 1). Although the sequence length (1.2 Gb) was 26.7% shorter than the expected size (4.5 Gb), 95.4% of complete BUSCOs were represented in the contigs (Table 1).

**Table 1.**
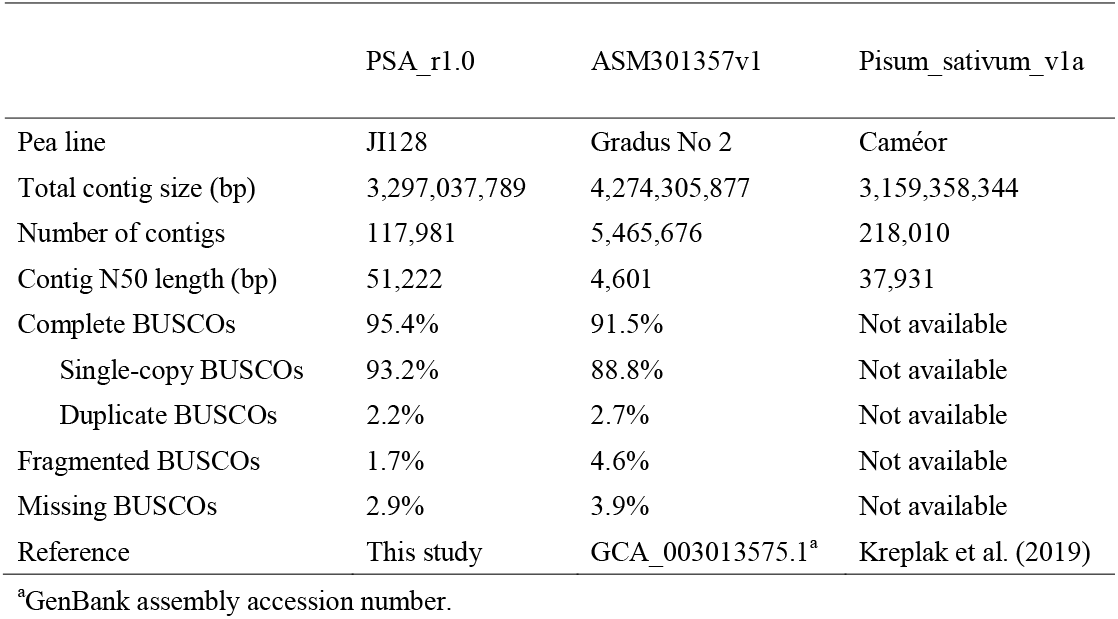
Statistics of the genome assemblies of pea lines.

**Figure 1.**
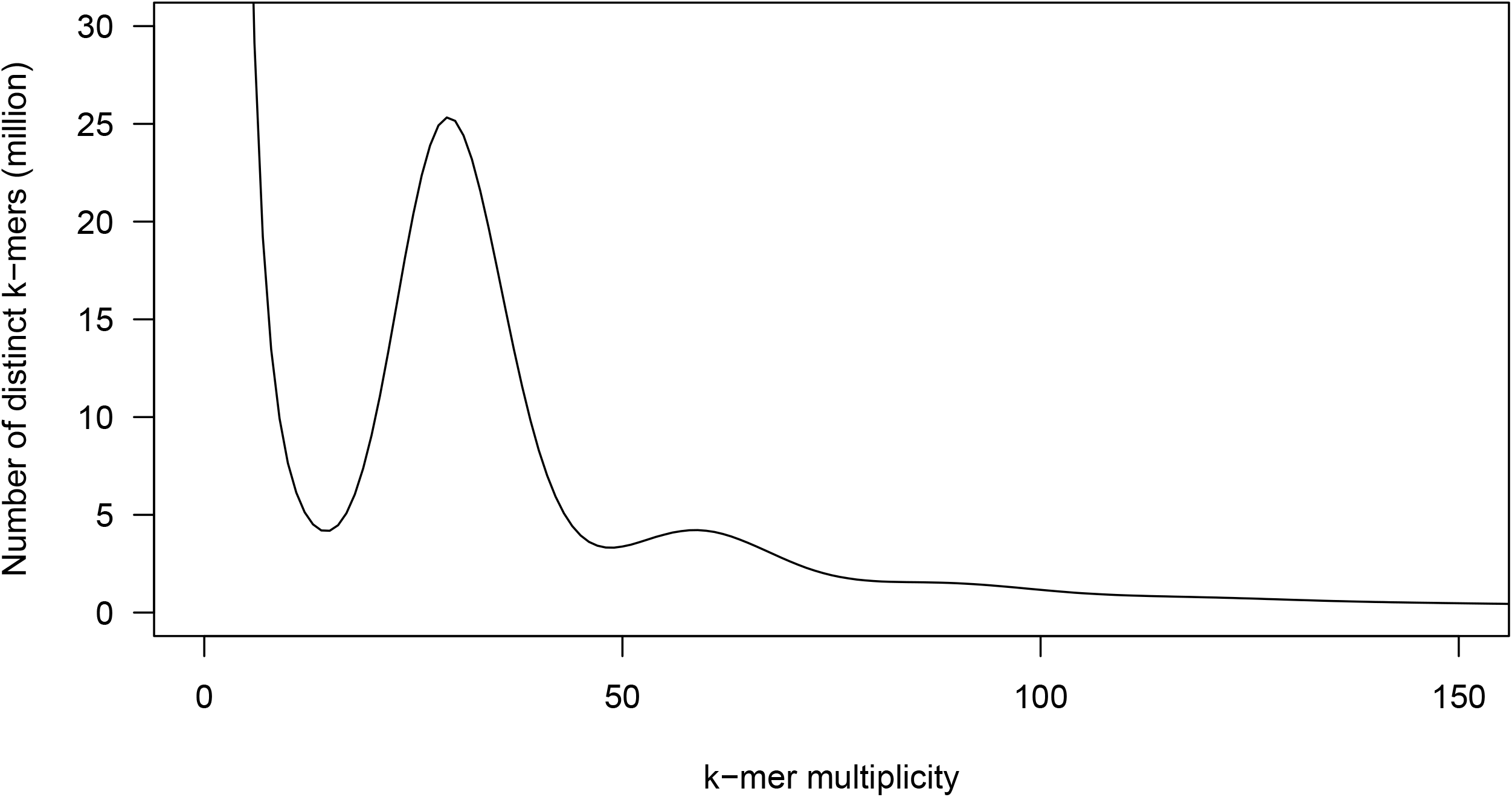
Genome size estimation for the pea line, JI128, based on the distribution of the number of distinct *k*-mers (*k*□= □17), with the given multiplicity values.

Short-read data of JI128 (80.3 Gb) and JI4 (68.7 Gb) obtained using the Illumina NextSeq 500 system (Table S1) were mapped onto the PSA_r1.0, Pisum_sativum_v1a, and ASM301357v1 assemblies to detect 6,374,054, 6,048,006, and 6,152,891 high-confidence SNPs, respectively, between the two lines were selected.

### Genetic map construction

A genetic map was constructed based on SNPs identified by ddRAD-Seq analysis of the F2 population, F1 hybrid, and parental lines. A total of 2.4 M reads were obtained per sample using ddRAD-Seq (Table S1), of which 95.1% were mapped onto the PSA_r1.0 assembly. High-confidence SNPs were selected from 2,183 loci located on 939 contigs. The ddRAD-Seq reads were mapped in parallel onto two publicly available pea genome sequence assemblies, ASM301357v1 and Pisum_sativum_v1a, with mapping rates of 95.3% and 95.4%, respectively, leading to the identification of 2,304 and 2,246 SNPs, respectively. Out of a total of 6,733 SNPs (= 2,183 + 2,304 + 2,246), 6,238 were separated into seven linkage groups and ordered. The linkage groups were named according to Kreplak et al. (2019). The resultant 891.8 cM genetic map contained 6,023 SNPs assigned to 832 genetic bins (Figure 2, Table 2, Table S2), these SNPs mapped to 1,998, 2,034, and 1,991 loci (727, 740, and 726 bins, respectively) on the PSA_r1.0, ASM301357v1, and Pisum_sativum_v1a reference genome assemblies, respectively. While 641 bins were common to all three reference genome assemblies, 56, 33, and 23 bins were unique to PSA_r1.0, ASM301357v1, and Pisum_sativum_v1a, respectively (Figure 3).

**Table 2.** Genetic map length and number of single nucleotide polymorphisms (SNPs) and genetic bins.

| Linkage group | Integrated |  |  | PSA_r1.0 |  |  | ASM301357v1 |  |  | Pisum_sativum_v1a |  |  |
| --- | --- | --- | --- | --- | --- | --- | --- | --- | --- | --- | --- | --- |
|  | No. of SNPs | No. of genetic bins | Map length (cM) | No. of SNPs | No. of genetic bins | Map length (cM) | No. of SNPs | No. of genetic bins | Map length (cM) | No. of SNPs | No. of genetic bins | Map length (cM) |
| chr1LG6 | 576 | 89 | 105.4 | 197 | 80 | 94.8 | 191 | 79 | 105.4 | 188 | 74 | 94.8 |
| chr2LG1 | 758 | 104 | 101.9 | 257 | 95 | 101.3 | 255 | 92 | 101.9 | 246 | 87 | 101.9 |
| chr3LG5 | 788 | 109 | 133.2 | 258 | 95 | 128.7 | 262 | 98 | 133.2 | 268 | 99 | 128.7 |
| chr4LG4 | 774 | 116 | 132.2 | 264 | 104 | 131.9 | 252 | 100 | 132.2 | 258 | 98 | 131.9 |
| chr5LG3 | 1,257 | 151 | 152.3 | 413 | 128 | 152.3 | 434 | 128 | 152.3 | 410 | 137 | 152.3 |
| chr6LG2 | 1,020 | 135 | 122.4 | 330 | 115 | 121.2 | 350 | 127 | 122.4 | 340 | 119 | 122.4 |
| chr7LG7 | 850 | 128 | 144.4 | 279 | 110 | 144.4 | 290 | 116 | 144.4 | 281 | 112 | 144.4 |
| **Total** | **6,023** | **832** | **891.8** | **1,998** | **727** | **874.6** | **2,034** | **740** | **891.8** | **1,991** | **726** | **876.4** |

**Figure 2.**
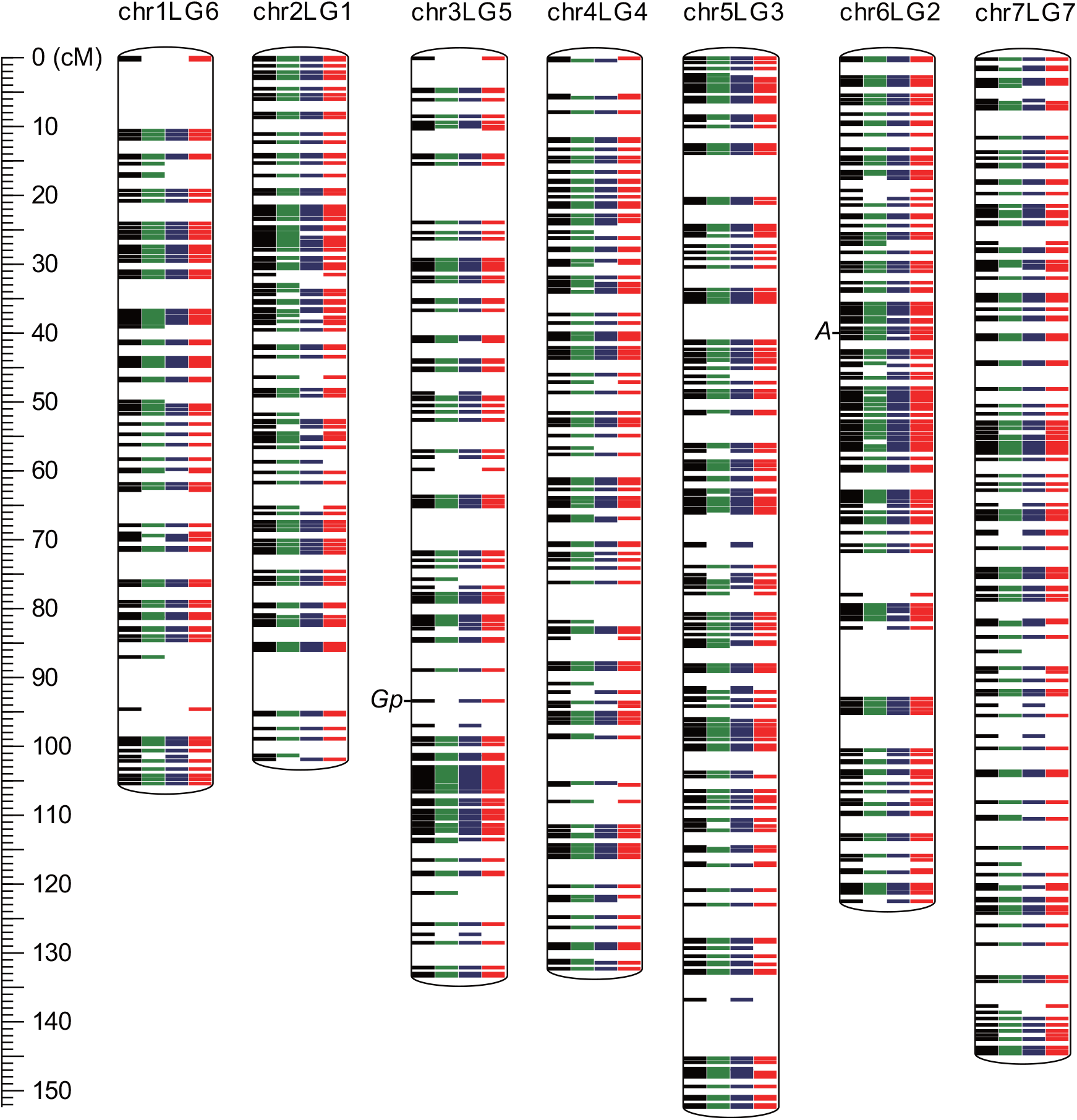
Genetic linkage map of pea. Green, blue, and red colors on chromosomes indicate genetic bins detected in the PSA_r1.0, Pisum_sativum_v1a, and ASM301357v1 genome assemblies, respectively, and black bars indicate consensus bins across the three assemblies. The genetic loci flower color and pod color are indicated as *A* and *Gp*, respectively.

**Figure 3.**
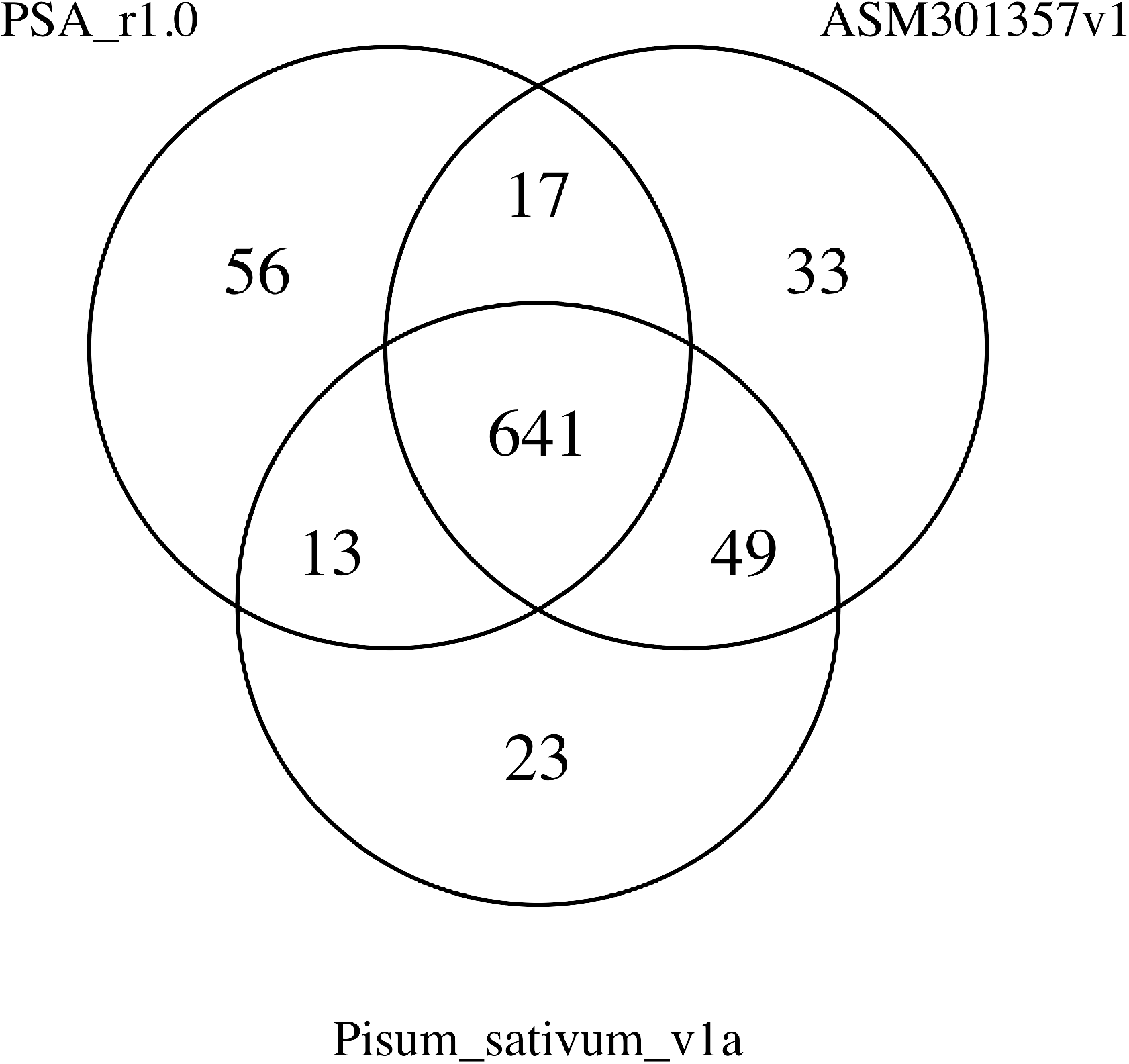
Number of genetic bins detected in the three reference genome assemblies, PSA_r1.0, Pisum_sativum_v1a, and ASM301357v1. The number of bins common or unique to the genome assemblies are shown by a Venn diagram.

On the basis of the genetic map, 918 contig sequences of PSA_r1.0 (135.8 Mb) and 1,001 contigs of ASM301357v1 (31.2 Mb) were anchored to the chromosomes (Supplementary Table S2). In Pisum_sativum_v1a, 91 scaffold and super-scaffold sequences (19.0 Mb) that had not been integrated into the seven pseudomolecule sequences (chr1LG6 to chr7LG7; 3.2 Gb) were newly assigned to pea chromosomes.

### Flower color gene at the A locus

Of the 167 plants in the F2 population, 109 produced red flowers and 44 produced white flowers (Figure 1); the remaining 14 plants were not evaluated for the flower color. Thus, the white:red segregation ratio fit the Mendelian segregation ratio for a single gene (*p* = 0.283; χ^2^ = 1.153). In the F1 progeny, red flower color was dominant to white flower color.

The flower color trait was mapped to a single locus on chr6LG2 (39.8 cM) of the genetic map (Figure 2). This mapping result was supported by GWAS (*p* = 2.9× 10^−77^). The physical location of this locus corresponded to 25,106 bp on Psa1.0_019194.1 (PSA_r1.0), 1,423 bp on PUCA013577394.1 (ASM301357v1), and 68,269,988 bp on chr6LG2 (Pisum_sativum_v1a). The SNP responsible for flower color variation in pea was located ∼60 kb proximal (or distal, whichever is correct) to the Psat6g060480 gene, which encodes a basic helix-loop-helix (bHLH) protein (Hellens et al., 2010). The SNP (A in JI4 and G in JI128) which has been proposed to cause the flower color variation was found at the splice donor site of the 6^th^ intron (at 68,336,837 bp on chr6LG2), based on the whole genome resequencing data of JI4 and JI128.

### Pod color gene candidates at the Gp locus

All F1 plants produced green pods. However, in the F2 population, 126 plants produced green pods, while 38 plants produced yellow pods (Figure 1); three lines had no pods at the time of phenotyping and therefore could not be evaluated for pod color. Thus, the pod color ratio fit the Mendelian segregation ratio for a single gene (*p* = 0.589; χ^2^ = 0.293). Together, these data indicate that pea pod color is controlled by a single gene, and green pod color is dominant to yellow pod color.

The pod color trait was genetically mapped to a single locus on chr3LG5 at 93.3 cM (Figure 2). This map position was close to the Gp locus studied by Mendel. The GWAS approach detected this position (*p* = 2.9 × 10^−117^) at 17,394 bp on scaffold04355 (PSA_r1.0) and at 1,901 bp on PUCA010639216.1 (ASM301357v1); both these physical positions are unknown in the pea genome.

While the sequence length of PUCA010639216.1 was only 5,818 bp, that of scaffold04355 was 64,468 bp, and three genes were predicted on scaffold04355, including Psat0s4355g0040, Psat0s4355g0080, and Psat0s4355g0120 (Figure 4). Psat0s4355g0040 was annotated as an unknown gene, while Psat0s4355g0080 and Psat0s4355g0120 were predicted as members of the 3’ exoribonuclease gene family. Interestingly, Psat0s4355g0080 and Psat0s4355g0120 were arranged head-to-head, with an interval of approximately 1.8 kb. Since these genes were potential candidate genes controlling the pod color trait, comparative genome and transcriptome analyses of JI4 and JI128 were performed.

**Figure 4.**
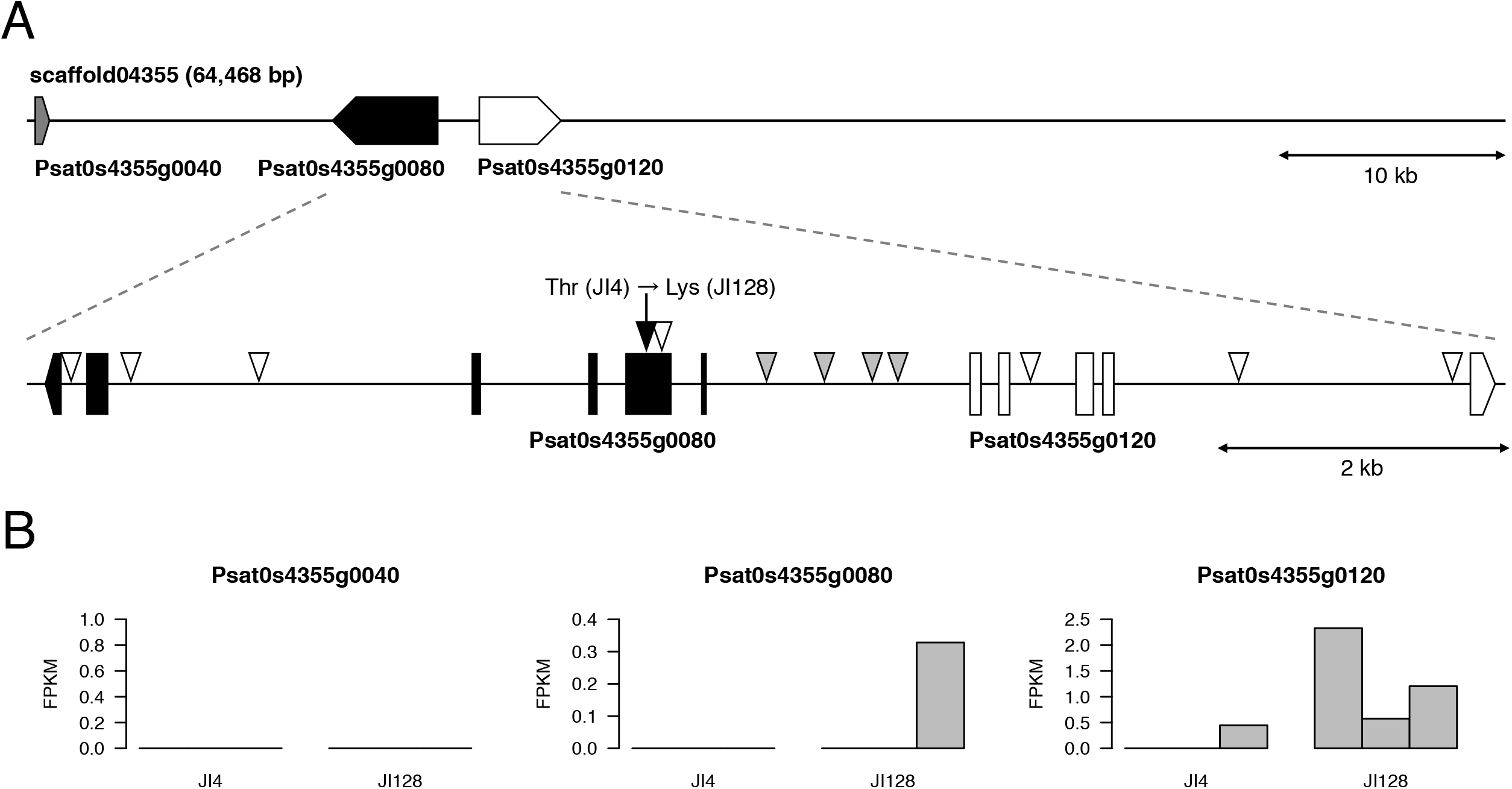
Genomic and transcriptomic analyses of scaffold04355 of Pisum_sativum_v1a. A. Genome structure of scaffold04355. Gray, black, and white boxes represent the Psat0s4355g0040, Psat0s4355g0080, and Psat0s4355g0120 genes, respectively. Triangles represent sequence variations between JI128 and JI4; white, black, and gray triangles represent silent, non-synonymous, and regulatory variants, respectively. B. Transcriptome-based RNA expression patterns of genes in scaffold04355. Bars indicate the expression levels of genes in three replicates of immature pods of JI128 and JI4, expressed as fragments per kilobase of exon model per million mapped reads (FPKM).

Whole genome resequencing analysis revealed a total of 36 sequence variations (28 SNPs and eight indels) between JI4 and JI128 on scaffold04355. Of these, only one SNP was a non-synonymous mutation, while the remaining 35 polymorphisms were silent mutations (six intronic variants, 28 intergenic variants, and one synonymous mutation) (Figure 4). Among the intergenic variants, two SNPs and two indels were located on the region between Psat0s4355g0080 and Psat0s4355g0120, which probably corresponded to the promoter regions of the two genes (Figure 4).

To analyze gene expression, three replicates of immature pods of each line were investigated by RNA-Seq. Approximately 21.2 M high-quality reads were obtained from each sample (Table S1) and mapped onto the Pisum_sativum_v1a reference genome assembly with a mapping rate of 93.9%. Out of 44,756 genes predicted in Pisum_sativum_v1a, 25,726 genes were expressed in one of the six samples at least. The expression level of Psat0s4355g0120 in JI128 (1.4) was approximately 10-fold higher than that in JI4 (0.15) (Figure 4). No obvious differences were detected in the expression levels of Psat0s4355g0040 and Psat0s4355g0080 between the two pea lines.

## DISCUSSION

In this study, we report the genome sequence assembly of the pea line, JI128, possessing yellow pods. Contigs were genetically anchored to the publicly available pea reference genome assemblies, Pisum_sativum_v1a and ASM301357v1. Our analysis provides information on the common genetic loci across the different genome sequence assemblies. Furthermore, the locus associated with pod color was identified on the genetic map, and the genes underlying this locus were identified on the reference genome sequence assemblies. Genomic and transcriptomic analyses revealed candidate genes responsible for the differences in pod color between JI128 and JI4.

To construct the genome sequence assembly of JI128 (PSA_r1.0), we employed a long-read sequencing technology of Oxford Nanopore Technologies (44× genome coverage). The PSA_r1.0 assembly showed higher contiguity (N50 = 51,222 bp) than the ASM301357v1 assembly (N50 = 4,601 bp), which was generated from Illumina short-read sequences of a single library (86× genome coverage), and similar to the Pisum_sativum_v1a assembly (N50 = 37,931 bp), which was constructed using Illumina short reads of multiple insert-size libraries (281× genome coverage) (Kreplak et al., 2019) (Table 1). However, the contiguity of assembled sequences obtained in this study was lower than expected. No contig sequences for the pod color locus were represented in the PSA_r1.0 assembly. It might be necessary to extend the sequences by integrating whole genome profiles of a bacterial artificial chromosome (BAC) library and/or an optical mapping technology of BioNano Genomics, as reported previously (Kreplak et al., 2019). Alternatively, a PacBio long-read technology, which produces high-quality long reads known as HiFi reads, might improve the assembly (Lang et al., 2020; Wenger et al., 2019).

The PSA_r1.0 assembly, together with contigs of ASM301357v1, was anchored to the pseudomolecule sequences of Pisum_sativum_v1a (Table S2). In the ddRAD-Seq analysis, 6,023 SNP loci were identified on the genetic map, and 918 and 1,001 contigs of PSA_r1.0 and ASM301357v1, respectively, were genetically assigned to the pseudomolecule sequences of Pisum_sativum_v1a, despite the short lengths of the anchored sequences (PSA_r1.0: 135.8 Mb; ASM301357v1: 31.2 Mb), because of the low contiguity of the assemblies. Thus, it is important to improve the assembly contiguity and/or increase the number of genetic loci to cover most of the contig sequences. Whole genome resequencing of mapping populations (Mascher et al., 2013) has become a realistic alternative to ddRAD-Seq, owing to the reduction in the cost of sequencing. The genetic anchoring approach would be useful for genetics in the pan-genome era (Bayer et al., 2020).

The 3’ exoribonuclease protein encoding genes, Psat0s4355g0080 and Psat0s4355g0120, were identified as potential candidates for the pod color. Psat0s4355g0120 was expressed in JI128, while Psat0s4355g0080 showed a missense mutation. Therefore, both these genes are currently considered as potential candidates for pod color. While the 3’ exoribonuclease proteins are reported to involve mRNA degradation in Arabidopsis (Nguyen et al., 2015), the function of these genes was not analyzed in pea. We propose a hypothetical scenario in which these genes lead to the formation of yellow pods. The 3’ exoribonuclease family proteins might suppress the production of a green pigment, probably chlorophyll, in pods or degrade the products, for example, in the green cotyledons of pea (Armstead et al., 2007; Sato et al., 2007). However, while the yellow color was dominant to the green color in cotyledons, an opposite trend was observed in pods. Since the 3’ exoribonuclease family proteins form dimers (Zuo and Deutscher, 2001), this dimerization might be required to suppress the factors involved in the biosynthesis of the green pigment. In heterozygous plants, if the protein dimer contained one subunit encoded by the dominant allele and the other subunit encoded by the recessive allele, the resulting dimer might lose function in accordance with the dominant-negative effect (Veitia, 2007), which would explain the results obtained in this study. However, further investigation is needed to verify this hypothesis and to identify the gene for the green pod color.

In conclusion, we determined the genome sequence of a pea line, JI128, possessing yellow pods, and identified the genetic locus responsible for the pea pod color. Since the map position of the pod color locus was closed to the locus reported by Mendel, we predict that the identified genomic region contains the gene responsible for the pod color of pea. Thus, the results of this study facilitate the identification of the gene controlling one of the seven traits studied by Mendel, and are expected to facilitate pan-genome studies in pea.

## ACCESSION NUMBERS

The sequence reads were deposited to the Sequence Read Archive (SRA) database of the DNA Data Bank of Japan (DDBJ) under the accession numbers DRA010800 and DRA010801. The DDBJ accession numbers of the assembled contig sequences, PSA_r1.0, are BNEU01000001-BNEU01117981.

## ACKNOWLEDGMENTS

We thank Dr. M. Ambrose in John Innes Center (Norwich, UK) for providing the seeds of pea lines JI128 and JI4. We also thank S. Sasamoto, S. Nakayama, A. Watanabe, Y. Kishida, C. Minami, and H. Tsuruoka at the Kazusa DNA Research Institute (Chiba, Japan) for technical assistance, and the technical staff members at Department of Technical Development in the Institute for Sustainable Agro[ecosystem Services (ISAS; The University of Tokyo, Japan) for the maintenance of plant material. This work was supported by the Kazusa DNA Research Institute Foundation.

## SUPPORTING INFORMATION

**Table S1.** Data obtained by whole genome resequencing, RNA-Seq, and ddRAD-Seq analyses of pea lines, JI128 and JI4.

**Table S2.** Genetic map and anchored genome sequences of JI128.

